# Extracellular rRNA profiling reveals the sinking and cell lysis dynamics of marine microeukaryotes

**DOI:** 10.1101/2024.05.31.596594

**Authors:** Hisashi Endo, Yuki Yamagishi, Thi Tuyen Nguyen, Hiroyuki Ogata

**Affiliations:** Bioinformatics Center, Institute for Chemical Research, Kyoto University, Gokasho, Uji, Kyoto, 611-0011, Japan; Shimadzu Techno-Research, Inc., Nishinokyo-Shimoaicho, Nakagyo-ku, Kyoto, 604-8436, Japan

**Keywords:** Phytoplankton, Protists, Cell-free RNA, Marine ecosystems, Biogeochemical cycles, Mesopelagic ocean

## Abstract

Marine plankton communities consist of numerous species, and their composition and physiological states are closely linked to ecosystem functions. Understanding biogeochemical cycles requires measuring taxon-specific mortality due to viral lysis or sloppy feeding, as the dissolved organic matter released contributes to rapid nutrient recycling and long-term carbon sequestration following microbial transformation. This study introduces a quantitative and comprehensive analysis of microeukaryotes in the dissolved constituents of seawater by using Mortality by Ribosomal Sequencing (MoRS) method. Our experimental pipeline successfully recovered 83% of cell-free rRNA. The ratio of cell-free rRNA to cell-associated rRNA was more than 10-fold higher in the mesopelagic layer than in the upper epipelagic layer, suggesting the mesopelagic zone as a hotspot for eukaryotic cell lysis. Many protist lineages, including phytoplankton such as haptophytes, are less susceptible to cell lysis in the epipelagic layer yet are actively lysed in the mesopelagic zone. Notably, over 86% of the significantly lysed species in the mesopelagic layer showed a habitat preference for the epipelagic layer. These findings indicate that sinking from the surface and lysis in the mesopelagic are prevalent dynamics for various eukaryotes.

## 1. Introduction

Protists, a diverse group of microscopic eukaryotes, are crucial components of marine ecosystems and biogeochemical cycles, playing roles in food webs, nutrient cycling, and carbon sequestration^1^. Studies on marine ecosystems and biogeochemical cycles have focused on specific lineages, such as diatoms and calcifying haptophytes, which can be distinguished by their microscopic morphology and chemotaxonomic pigment features. Recent studies based on high-throughput sequencing (HTS) have underscored the diversity of eukaryotic plankton, including protozooplankton, and the remarkable contributions of these species to natural assemblages^2,3^. Therefore, elucidating the complex interactions between diverse microorganisms is conducive to understanding marine systems^4,5^.

Viruses and multicellular predatory organisms are also important members of marine microbial consortia as they control the abundance and composition of protists via lytic infection and grazing. The plankton mortality due to viral lysis has significant consequences for marine biogeochemistry. The lytic processes cause cell contents to be released into the surrounding seawater as dissolved organic matter (DOM; defined as organic matter passing through a 0.2–0.7 µm filter). According to field experiments, viral infection eliminates 10–40% of plankton populations daily^6–8^. Besides, sloppy feeding by predators yields similar effects, as it converts large amounts of cell content to DOM^9,10^, whereas an engulfment process like phagocytosis incorporates whole prey cells into the biomass of higher trophic levels^11^. Incubation experiments indicated that sloppy feeding of copepods on phytoplankton cells released 10–36% of cellular organic carbon as DOM^10^. These processes causing cell lysis (i.e., breakdown of cell membrane and the release of intracellular contents) can boost nutrient recycling on the surface water at the expense of reducing sinking carbon export to the deep ocean (biological carbon pump)^6,12^. On the other hand, cell lysis can occur during the sinking processes via viral infection and deep-sea predators, which results in remineralization in the deep ocean^13–15^. However, the relationship between sinking processes and the timing of cell lysis remains poorly understood.

Cell-free nucleic acids (DNA and RNA) are key DOM products released by cell lysis processes^16^. These nucleic acids exist in viral particles, extracellular vesicles, and free nucleotide molecules resulting from cell breakdown or excretion. RNA molecules comprise a major form of DOM that accounts for 60% of the dissolved organic phosphorus (DOP) in seawater, and their concentration exceeds that of DNA by up to 3– 10 times^17–19^. Nucleic acids are seldom excreted spontaneously from living cells due to their high content of valuable elements such as nitrogen and phosphorus (C:N:P = 10:4:1)^16,20^. The high bioavailability and low structural stability render cf-RNA a labile DOM with a short half-life (a few hours to several days)^21^. These considerations imply that RNA in the DOM pool is continuously supplied from damaged and burst cells. Therefore, it provides information on recent or ongoing cell lysis events caused by viral infection and sloppy feeding by zooplankton.

A recent milestone study introduced the concept of Mortality by Ribosomal Sequencing (MoRS), which evaluates cell lysis by analyzing ribosomal RNA sequences in the extracellular fraction^22^. They demonstrated that extracellular rRNA (synonymous with cell-free rRNA; cf-rRNA) is actively produced by viral lysis and can thus serve as a proxy for evaluating the top-down control of prokaryotic populations. Remarkably, when combined with HTS, this molecular marker allows for the simultaneous evaluation of the degree of viral lysis for hundreds to thousands of microbial species in an environment without information on virus-host relationships^22,23^. Although the lytic processes of microeukaryotes would be more complicated considering the multiple factors, including viruses, sloppy feeding, and other mechanical stresses, cf-rRNA would provide valuable insights into the mortality of eukaryotes.

In this study, we applied the concept of MoRS to evaluate the cell lysis of eukaryotes. Nevertheless, extracting sufficient RNA for quantification/sequencing from natural seawater filtrates remains methodologically challenging, especially in oligotrophic and aphotic environments. Concentration-based methods, such as tangential flow filtration and ultracentrifugation, are typically used to investigate cell-free nucleic acids in seawater^24,25^. These techniques require considerable time and effort, and the RNA is biased toward a specific molecular size or degraded during handling. We established a method for extracting cell-free RNA from a large volume (> 40 mL) of dissolved seawater (filtrate of 0.2 µm pore filter) without concentration treatment. This method recovered > 80% of cf-rRNA, yielding sufficient rRNA pools for abundance and taxonomic analyses of eukaryotes, even from oligotrophic subtropical surface water and deep water characterized by a low density of plankton cells. This method allowed us to obtain comprehensive profiles of cell lysis of eukaryotic plankton based on cell-associated (living) and cell-free (dead) fractions of rRNA quantification and metabarcoding (Fig. 1).

**Figure 1.**
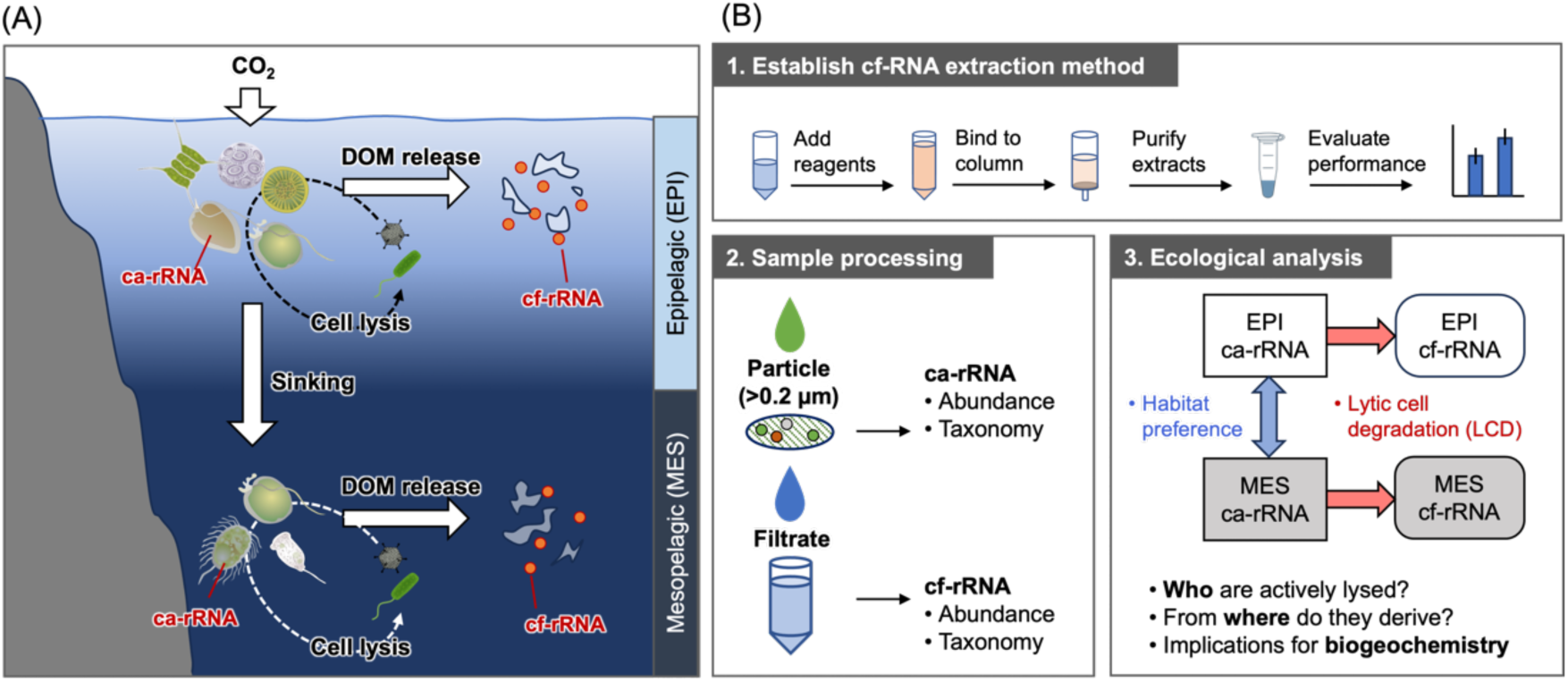
The concept of this study. (A) A schematic diagram showing the evaluation of taxon-specific mortality and vertical difference of microeukaryotes. (B) The workflow to evaluate the mortality of marine protists by cell-free 18S rRNA profiling. Abbreviations: EPI, Epipelagic (0–200 m depth); MES, Mesopelagic (200–1,000 m); DOM, Dissolved organic matter; ca-rRNA, Cell-associated rRNA; cf-rRNA, Cell-free rRNA.

## 2. Material and methods

### 2.1. Sample collection

Field sampling was conducted at five stations near the Kuroshio off south Kyushu, Japan, during the KS-22-15 cruise (October 16–27, 2022) of the *R/V Shinsei Maru* (JAMSTEC). Seawater samples were collected from the surface (10 m), deep chlorophyll *a* maximum (SCM, 35–93 m), and mesopelagic (300 m or 500 m) layers using Niskin bottles attached to a CTD-RMS system (Table S1).

Samples for the cell-associated and cell-free rRNA analyses were prefiltered with a 144 µm pore-size mesh to remove metazoa. Then seawater (300 mL) was filtered through the 47 mm diameter 0.22 µm pore size polycarbonate (PC) filter (Isopore GTTP04700, Merck Millipore) with a gentle vacuum (< 0.013 MPa) to separate into cell-associated and cell-free rRNA fractions. The filter samples (cell-associated RNA) were soaked in 600 µL buffer RLT (Qiagen) containing 1% *β*-mercaptoethanol and 0.2 g of muddled 0.1 mm glass beads. Forty milliliters of the filtrates (cell-free rRNA) were subsampled into 50 mL Falcon tubes (Watson). The filter and filtrate samples were flash-frozen in liquid nitrogen and stored in a deep freezer.

Seawater samples (500 mL) for light microscopic analysis were preserved with Lugol’s solution (1% v:v final concentration). The samples were concentrated by precipitation and examined using an upright microscope (BX50, Olympus).

### 2.2. Nucleic acid extraction and cDNA synthesis

Cell-associated RNA samples were extracted using the RNeasy Plus Mini Kit (Qiagen) following the manufacturer’s protocol, with some modifications. We used buffer RLT instead of buffer RLT Plus (Qiagen), which includes detergents, as the lysis buffer because it yielded a higher RNA concentration in a preliminary experiment. The ice-cold lysis buffer was agitated with a bead beater at 2,500 rpm thrice for 50 s before starting the protocol.

To extract cell-free RNA from the seawater filtrate samples (< 0.22 µm), we used the Wizard Enviro Total Nucleic Acid (TNA) Kit (Promega), according to the manufacturer’s instructions with an elution volume of 40 µL. This kit was developed to capture and concentrate viral TNA from a large volume of wastewater (> 40 mL) using a column-based aspiration system^26^.

Genomic DNA removal and first-strand cDNA synthesis were conducted for the extracted nucleotides using SuperScript IV VILO Master Mix with the ezDNase enzyme kit (Thermo Fisher Scientific) following the manufacturer’s specifications.

### 2.3. Performance evaluation of cell-free RNA extraction method

We conducted an RNA spike-in extraction experiment to assess the extraction efficiency of cell-free (dissolved) rRNA using a Wizard Enviro TNA Kit. Purified ribosomes from *Escherichia coli* strain B (New England Biosciences, cat. P0763S) were suspended in artificial seawater, and the solution was subjected to TNA extraction and cDNA synthesis using the abovementioned procedure. For comparison, the same ribosomal solution was used directly for cDNA synthesis. We also evaluated the performance of the ca-rRNA extraction protocol using the RNeasy Plus Mini Kit (Qiagen). All experiments were performed in triplicate.

The concentration of cDNA synthesized from *E. coli* 16S rRNA was quantified using a digital PCR system (QIAcuity One 5plex, Qiagen) in a Nanoplate 8.5K 96-well plate. A specific primer pair, 16S_EC442F and 16S_EC636R, was designed and used in the QIAcuity EG PCR Kit (Qiagen) following the manufacturer’s instructions (Table S2).

Additionally, we tested the adsorption properties of the PC filter (Isopore GTTP04700, 47 mm diameter, 0.22 µm pore size) to confirm that the filter membrane did not adsorb ribosomes. For comparison, we evaluated two other membrane filters made of polyvinylidene difluoride (PVDF) (Durapore GVWP04700) and mixed cellulose esters (MCE) (MF-Millipore GSWP04700), which have the same mean pore sizes but different chemical composition and structural conformations. Artificial seawater, including the purified ribosomes from *E. coli* strain B, was filtered through each filter in triplicate, and the filtrates were subjected to TNA extraction and cDNA synthesis. The absorption of ribosomes to each filter was measured by quantifying *E. coli* 16S rRNA cDNA using the abovementioned method.

### 2.4. Droplet digital PCR

A QX200 Droplet Digital PCR (ddPCR) system (Bio-Rad) was used to quantify eukaryotic rRNA. This system enables absolute quantification of the target sequence without a standard curve by counting the nucleic acid molecules encapsulated in discrete nanodroplets. We used the primer pair E572F and E1009R to target the V4 region of the small subunit of the 18S rRNA gene (Table S2)^27^

The 20 µL reaction mixtures containing primers (0.4 µM in final concentration), the cDNA template, and the ddPCR EvaGreen Supermix (2X; Bio-Rad) were prepared and then loaded into a DG8 Cartridge, followed by 70 µL of Droplet generation Oil. Droplets produced using a droplet generator (Bio-Rad) were transferred to 96-well PCR plates. The amplifications were performed using a thermal cycler with the following conditions: 95 °C for 5 min, denaturation at 95 °C for 30 s, and annealing/extension at 54 °C for 2 min for 40 cycles. After amplification, the plate was analyzed using a QX200 Droplet Reader (Bio-Rad).

### 2.5. SSU rRNA metabarcoding

Amplicon sequencing was performed as described previously^28^. Briefly, the V4 region of the 18S rRNA gene was amplified in triplicate from the cDNA template using the universal eukaryotic primer set E572F/E1009R (Comeau et al., 2011) attached to Illumina overhang adapters using KAPA HiFi HotStart ReadyMix. The thermal cycling condition was as follows: a pre-denaturation step at 98 °C for 30 s, followed by 30 cycles of denaturation at 98 °C for 10 s, annealing at 61 °C for 30 s, and extension at 72 °C for 30 s, with a final extension at 72 °C for 5 min. Triplicate PCR products of approximately 440 bp in length were mixed after purification using Agencourt AMPure XP beads (Beckman Coulter). Amplicon libraries were prepared using the Nextera XT Index Kit V2 and sequenced on an Illumina MiSeq platform using 300 bp paired-end sequencing.

### 2.6. Sequencing processing

Sequence data were analyzed using QIIME2 (ver. 2023.9)^29^ with the DADA2 plugin^30^. Before being imported into the pipeline, raw reads were processed using fastp^31^ to remove low-quality reads and adapter sequences. Primer sequences were removed from the quality-controlled reads with the “cutadapt trim-paired” command, then the paired-end reads were merged and further subjected to denoising, dereplication, and chimera removal to generate amplicon sequencing variants (ASVs) with the “dada2 denoise-paired” command. Rare ASVs that appeared in less than ten sequences across all samples or those found in only one sample were excluded. Taxonomy was assigned to remaining ASVs based on a pre-trained naive Bayes classifier (trained against the PR2 reference database version 4.14.0) using the “q2-feature-classifier” plugin^32^. ASVs classified as non-protist lineages (metazoans, prokaryotes, and unclassified) were excluded from the table. The read count of each sample was rarefied to 25,878 using random sampling.

### 2.7. Ecological analyses

To show the taxonomic composition of the microeukaryotes, a heatmap was created based on the log-transformed relative abundance of 18S rRNA at the class level (PR2_Lv5). Hierarchical clustering based on the Ward method with a Euclidean distance measure was performed to evaluate similarities across taxa. Samples were also clustered based on the Bray–Curtis dissimilarity of the ASV-level taxonomic composition (not at the class level) using Ward’s D2 method. Statistical differences in sample groups (depths, forms, and sample sites) were assessed for ASV abundance using an analysis of similarities (ANOSIM) with 9,999 permutations. All statistical analyses were conducted using the R package “vegan” (https://cran.ism.ac.jp/web/packages/vegan).

### 2.8. Analysis of DA ASVs

Differences in ASV abundance between the two sample categories were tested using edgeR (version 4.0.16)^33^ in R. ASVs that appeared in fewer than four samples were removed from the dataset. Read counts of each ASV were adjusted based on the trimmed mean of M-values (TMM) normalization factor and the negative binomial dispersion as estimated through the ‘calcNormFactors’ and ‘estimateDisp’ functions. We fit a generalized linear mode to normalize read counts using the ‘glmFit’ command after adding a small prior count of 0.125 to avoid undefined ASVs when either of the counts is zero. Then, statistics of differential abundance were assessed using the ‘glmLRT’ command. ASVs with a False Discovery Rate (FDR) < 0.1 and an absolute value of log_2_-fold change (logFC) > 2 were considered significantly differentially abundant (or less abundant) ASVs (DA ASVs).

We performed DA analysis on three sets of comparisons: (1) cf- and ca-rRNA forms in epipelagic samples, (2) cf- and ca-rRNA forms in mesopelagic samples, and (3) epipelagic and mesopelagic samples of ca-rRNA. The logFC values between the cf- and ca-rRNA forms were defined as the cell-lysis index (CLI), and those between the epipelagic and mesopelagic layers were defined as the epipelagic enrichment index (EEI).

### 2.9. Habitat distribution analysis

To test the bimodality of the EEI measure (i.e., whether a protist lineage consists of ASVs adapted to different depths), the function ‘Modes’ in the R package “LaplacesDemon” (https://github.com/LaplacesDemonR) was used. This function determines the number and value of local maxima in the Gaussian kernel density distribution of variables with at least 10% of the distribution area. We regarded a protist lineage (subdivision level, PR2_Lv4) with at least one mode in each depth category (epipelagic and mesopelagic zones) as a multi-habitat lineage.

## 3. Results

### 3.1. Feasibility of quantitative analysis of cell-free RNA in seawater

The extraction efficiency of RNA was 83.1 ± 6.5% (mean ± standard deviation, SD) for the cf-fraction and 21.9 ± 5.7% (mean ± SD) for the cell-associated fraction (n = 3; Fig. S1A) as measured by rRNA abundance. We confirmed the linearity of the extraction volumes from 2.0 × 10^2^ to 2.0 × 10^6^ copy mL^−1^ of rRNA dissolved in seawater (log-log linear regression: *r*^2^ = 0.995, *p* < 0.001, slope = 1.012, n = 15; Fig. S1B).

By testing the absorption of the ribosome to three different membrane filters, we confirmed that no filtration loss occurred for the PC and PVDF filters (pairwise *t*-test, p > 0.05, n = 3; Fig. S2). In contrast, nearly 80% of the 16S rRNA was lost after the filtration by the MCE filter (pairwise *t*-test, p < 0.05), indicating that the filter adsorbed a large proportion of ribosome. This study used a PC filter for all the cf-rRNA sampling.

### 3.2. Hydrographic conditions in the study sites

The study sites were located across the Kuroshio axis (Fig. 2A). Water mass structures were clearly distinguished between the southern (stations 3 and E1) and northern sites (stations 5, 6, and 8) using the T–S diagram; northern stations were characterized by lower surface temperature and salinity (Fig. 2B). Northern stations had relatively higher chlorophyll *a* fluorescence signals than southern stations (Fig. 2C).

**Figure 2.**
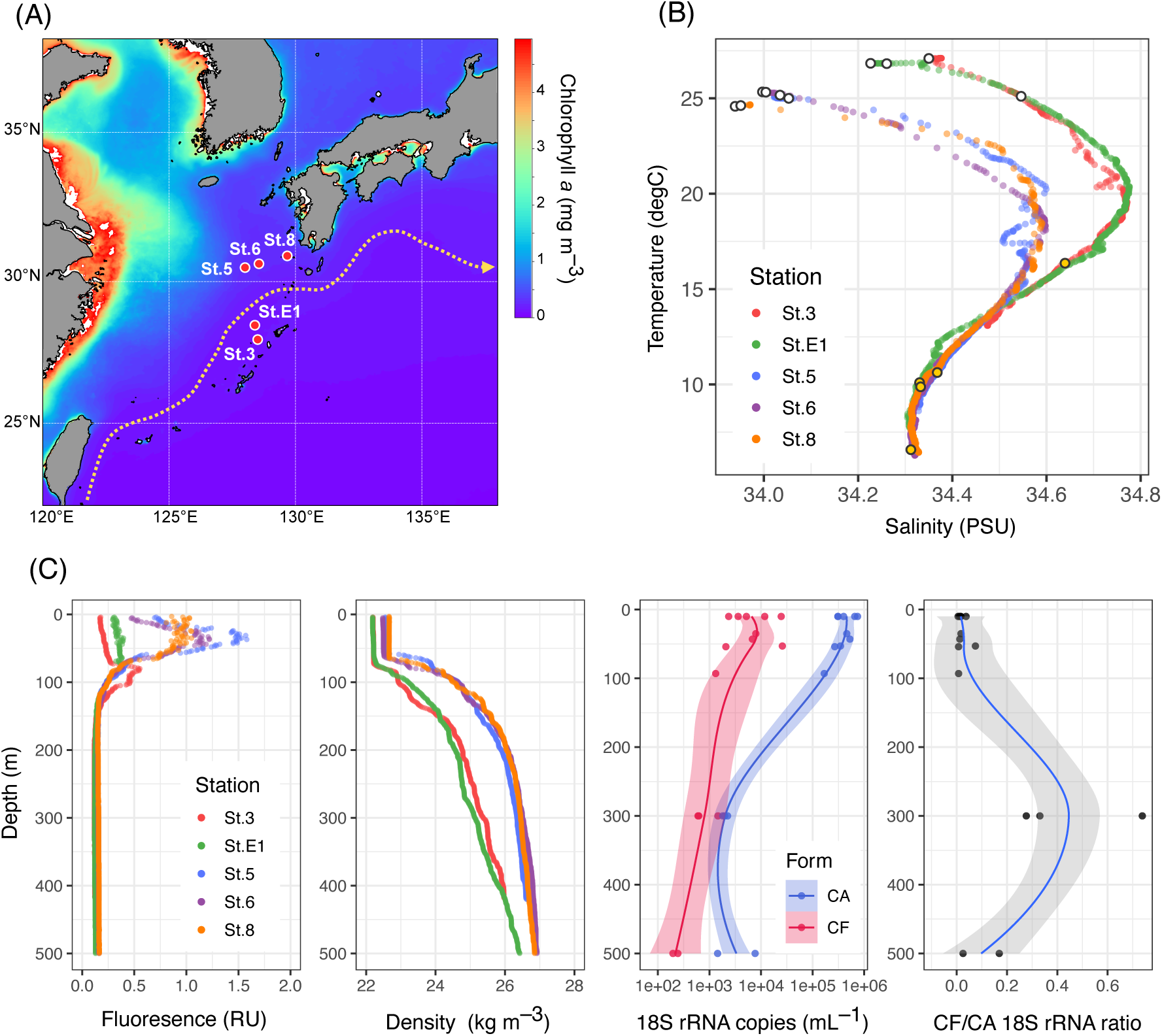
Location and environment of the study area. (A) A map of the sampling sites. The background image represents chlorophyll *a* concentration archived from MODIS-Aqua (monthly mean of October 2022, 4 km level 3 product). The dashed arrow indicates the position of the Kuroshio Current obtained from the Japan Coast Guard website (periods between October 10 and 17, 2022). (B) Temperature–salinity (T-S) diagram of all sampling sites. The sampling depths in the epipelagic and mesopelagic are indicated in the white and yellow circles, respectively. (C) Vertical profiles of fluorescence (relative unit of Seapoint chlorophyll fluorometer), seawater density (sigma-T), 18S rRNA abundance in different forms (cell-associated and cell-free fractions), and the abundance ratio of cell-free to cell-associated 18S rRNA (cf-rRNA/ca-rRNA).

### 3.3. Vertical abundance profiles of cell-associated and cell-free 18S rRNA

The abundance of 18S rRNA, as evidenced by ddPCR, varied across forms and depths (Fig. 2C). Cell-associated rRNA represented relatively high values in the euphotic (surface and SCM) layers (4.19 ± 1.80 × 10^5^ copies mL^−1^, mean ± SD, n = 10), but lower values in the mesopelagic layers (3.03 ± 2.62 × 10^3^ copies mL^−1^, n = 5). Cell-free rRNA showed higher values in the euphotic layers (9.126 ± 9.05 × 10^3^ copies mL^−1^) while lower values in the mesopelagic layers (6.23 ± 5.05 × 10^2^ copies mL^−1^). The ratio of the absolute abundances of cf-rRNA to ca-rRNA (cf-rRNA/ca-rRNA) ranged between 0.01–0.07 (0.02 on average) and 0.03–0.74 (0.31 on average) in euphotic and in mesopelagic layers, respectively.

### 3.4. Taxonomic composition of cell-associated and cell-free 18S rRNA

After removing rare and non-protist amplicon sequencing variants (ASVs), the sequencing analysis yielded 3,749 protist ASVs. The proportion of protist sequences to the total eukaryotic rRNA reads was 91.9% for cf-rRNA and 93.2% for ca-rRNA.

Based on the protist ASV composition, the original seawater samples were primarily clustered by sampling depth (surface epipelagic and deep mesopelagic zones; ANOSIM, R = 0.45, *p* < 0.01; Fig. 3). These groups were further categorized into cell-associated and cell-free rRNA forms (ANOSIM; R = 0.25, *p* < 0.01). Sampling locations were less important in explaining taxonomic composition differences (ANOSIM, R = 0.03, *p* = 0.21).

**Figure 3.**
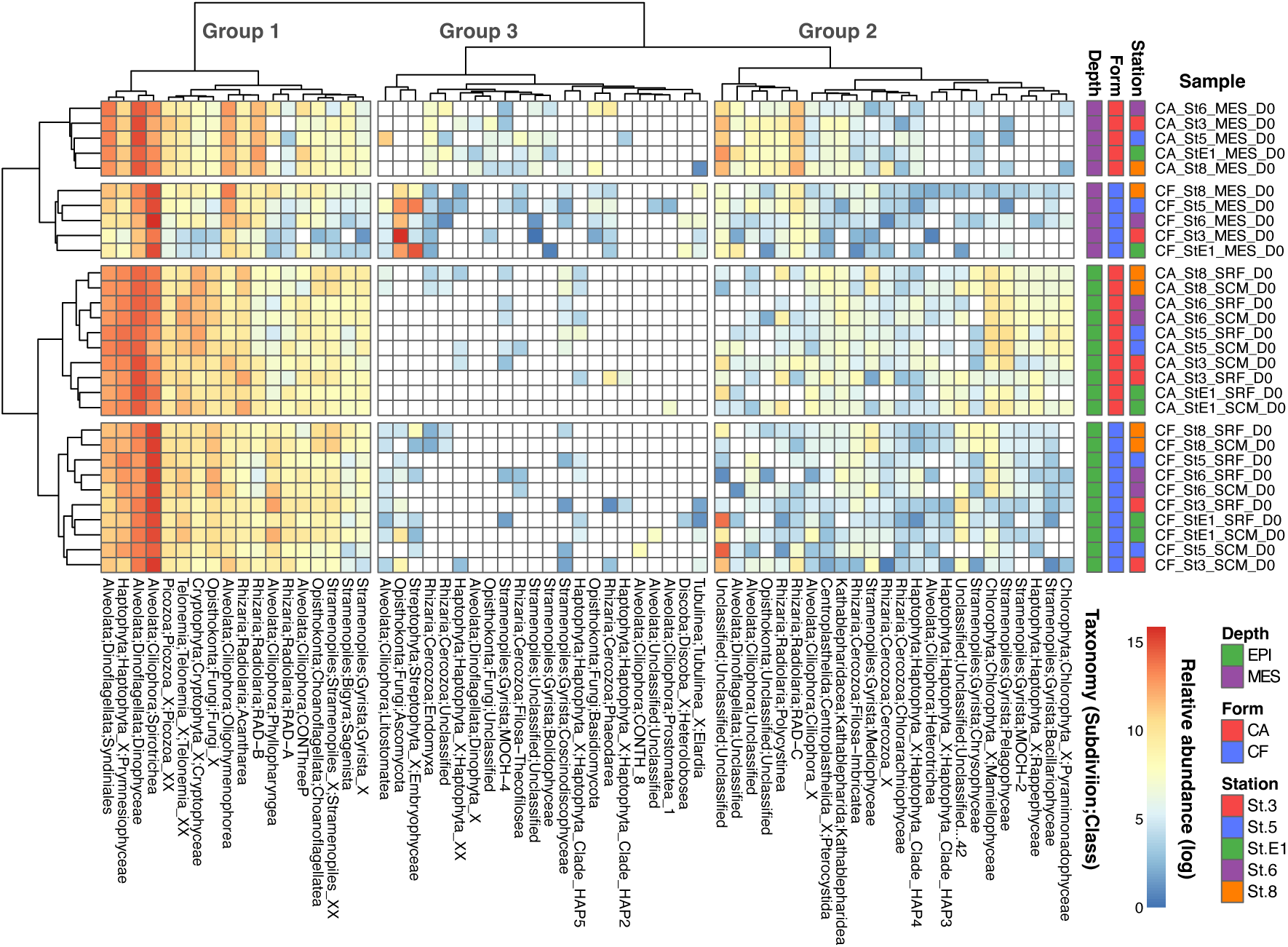
Heatmap and cluster analysis showing the taxonomic composition of protists. The heatmap indicates the log-transformed relative abundance of protist 18S rRNA at the class level (PR2_Lv5). Samples were clustered based on ASV-level taxonomic composition (not on the class level) in the left dendrogram. In the top dendrogram, protist clades were clustered based on their emerging patterns across samples at the class level. Protist classes with maximum contributions < 0.1% across all the samples were not shown.

Eukaryotic taxa were classified into three groups based on their emergence patterns (Fig. 3). Group 1 comprised ubiquitously distributed and relatively abundant lineages, such as Prymnesiophyceae (Haptophyta), Dinophyceae (Dinoflagellata), Spirotrichea (Ciliophora), Choanoflagellatea (Choanoflagellata), and Acantharea (Radiolaria). Group 2 was characterized by lineages with moderate contributions, including Mediophyceae, Bacillariophyceae, Chrysophyceae (Gyrista), and Mamiellophyceae (Chlorophyta). Group 3 was marked by lineages representing low relative abundance yet emerged endemically or sporadically in specific samples.

### 3.5. Differentially abundant (DA) ASVs in each rRNA form and habitat

After removing less prevalent ASVs, 1,082 ASVs were used for DA analysis. We detected 138 DA ASVs in epipelagic samples, 94 of which were significantly enriched in the cf-fraction (significantly lysed) and 44 in the ca-fraction (significantly intact; Fig. 4A–B). A total of 190 DA ASVs (171 lysed and 19 intact) were observed in mesopelagic samples. Vertical habitat comparison identified 661 DA ASVs; 437 and 224 were significantly enriched in upper epipelagic and deep mesopelagic samples, respectively.

**Figure 4.**
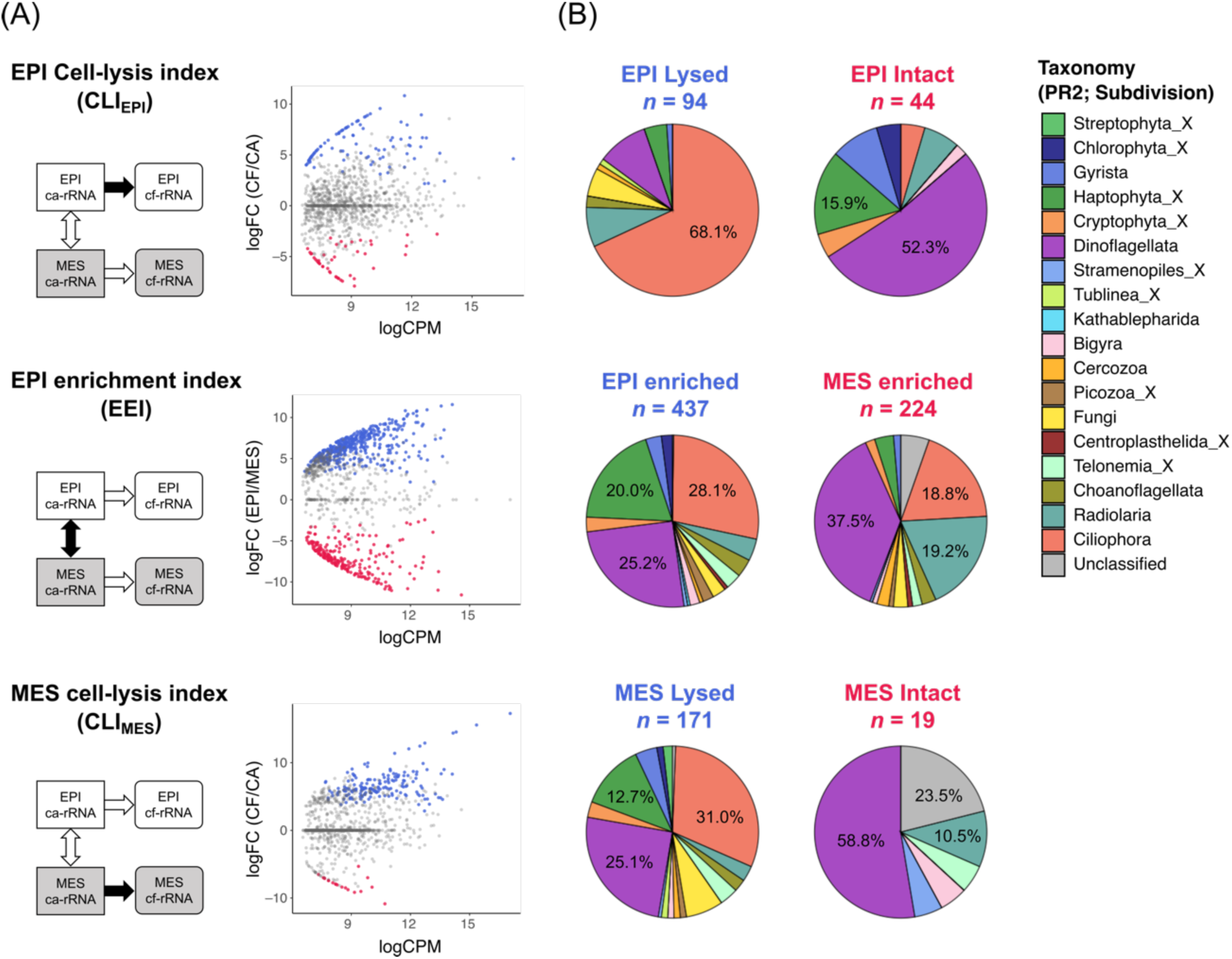
Differential abundance analysis assessing cell lysis and habitat characterization of protist communities. (A) MA-plot showing the average abundance (x-axis) and fold changes (y-axis) of all the ASVs. The upper and lower panels denote the differential abundances between cell-free and cell-associated fractions (i.e., cell-lysis index, CLI) in the epipelagic and mesopelagic, respectively. The middle panel indicates the abundance changes between epipelagic and mesopelagic communities (i.e., epipelagic enrichment index, EEI) of the cell-associated rRNA. (B) Pie charts summarizing the eukaryotic taxonomy (PR2_Lv4; subdivision level) with significant differences in each comparison.

The subdivision Ciliophora was the most abundant in terms of the number of lysed ASVs at both epipelagic and mesopelagic depths (Fig. 4B). Dinoflagellata and Haptophyta_X were the two most abundant lineages in intact ASVs in the epipelagic zone, although they were the second- and third-most lysed lineages in the mesopelagic zone.

### 3.6. Taxon specificity of prosperous and lytic depths

When comparing the ASVs that emerged in both the epipelagic and mesopelagic samples, 6 of the 12 eukaryotic subdivisions tested (Dinoflagellata, Ciliophora, Haptophyta_X, Gyrista, Choanoflagellata, and Cryptophyta_X) displayed significantly higher CLI (cell-lysis index) in mesopelagic samples (Wilcoxon rank-sum test, *p* < 0.01, n = 8–156; Fig. 5A). The mean and median CLI values were higher in the mesopelagic region for other lineages, although the difference was not statistically significant or could not be estimated. All 12 major lineages showed a larger number of lysed DA ASVs in the mesopelagic zone than in the epipelagic zone (Fig. 5B).

**Figure 5.**
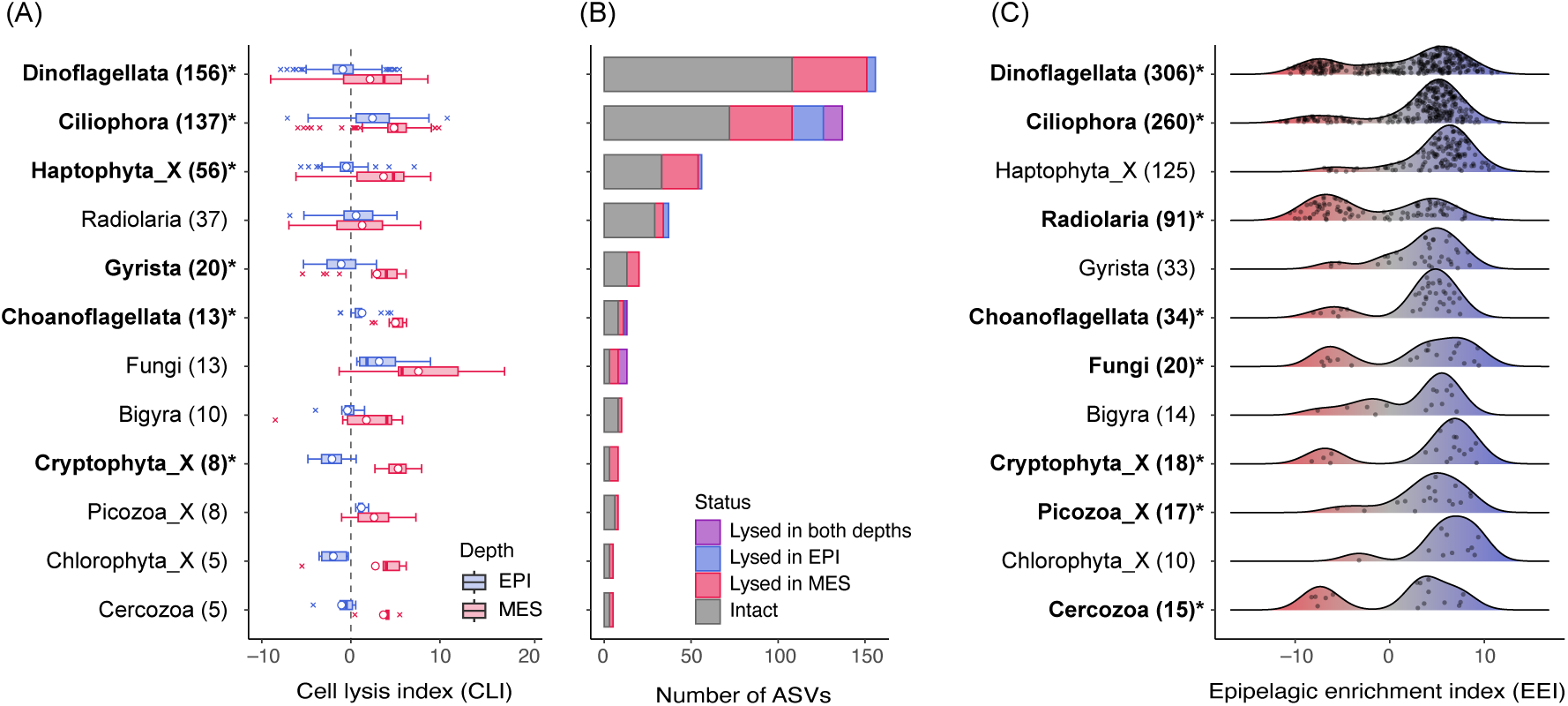
Taxon-specific lysis and habitat preference of protists. (A) Boxplot summarizing the lysis index of protist ASVs in each lineage (subdivision level) in the epipelagic (EPI) and mesopelagic (MES) zones (open circle, average; centerline, median; box limits, 25%–75% quantiles; whiskers, 1.5 × interquartile range). The *p-*value of the difference between EPI and MES < 0.01 is marked with an asterisk and bolded. (B) Number of ASVs and their lytic status of each lineage at each depth. (C) Density charts summarizing the epipelagic enrichment index (EEI) of protist ASVs in each clade. Values in the brackets indicate the number of ASVs in each lineage. A lineage having a bimodal distribution (i.e., comprising ASVs adapted to the epipelagic and mesopelagic zones) is marked with an asterisk and bolded. ASVs with zero values in either CLI (A and B) or EEI (C) measures were removed from all plots and statistics. Protist lineages having fewer than five ASVs were not shown.

The distribution of EEI values, a proxy for the preferred habitat depth, differed among eukaryotic groups. The protist lineages, which mainly consisted of phototrophic lineages such as Haptophyta_X, Gyrista (50% were diatoms), and Chlorophyta_X, showed unimodal density distributions with high ASV abundances in epipelagic layers (Fig. 5C). Conversely, heterotrophs such as Ciliophora and Radiolaria showed bimodal distributions, indicating that the lineages consisting of ASVs adapted to either mesopelagic or epipelagic habitats. Bimodal patterns were also observed for Dinoflagellata and Cryptophyta_X.

### 3.7. Characteristics of lysed eukaryotes in the deep sea

A significant positive correlation was detected between EEI (epipelagic enrichment index) and CLI_MES_ (CLI in the mesopelagic communities; Spearman’s rank test, *ρ* = 0.78, *p* < 0.001, n = 670; Fig. 6A). By comparing the DA ASVs of the EEI and CLI_MES_ categories, we found that 147 ASVs were shared between the two measures, corresponding to 86.0% of the DA ASVs in CLI_MES_ and 33.6% in EEI. The intersecting ASVs were mainly Ciliophora (29.9%), Dinoflagellata (27.9%), and Haptophyta_X (14.3%; Fig. 6B).

**Figure 6.**
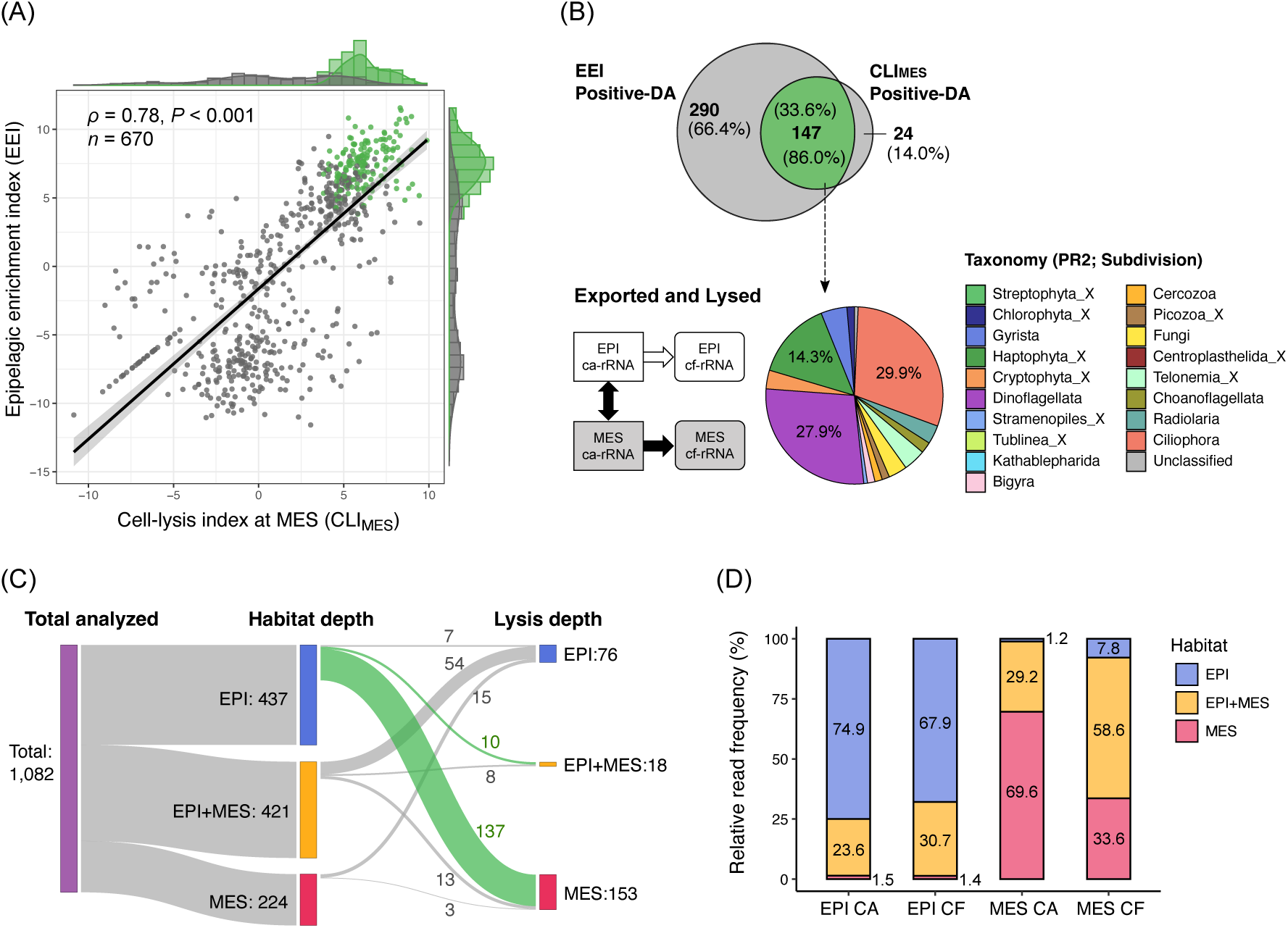
Detection of actively lysed microeukaryotes in the deep sea. (A) Relationship between the cell-lysis index in the mesopelagic communities (CLI_MES_) and the epipelagic enrichment index (EEI), a prosperity measure of sunlit habitats. Differentially abundant ASVs in both the CLI_MES_ and EEI are shown in green. (B) Inter-relationship and the taxonomic composition (subdivision level) of ASVs preferring the epipelagic habitat and actively lysed in the mesopelagic zone. (C) Sankey diagram displaying the number of ASVs classified into each habitat depth and lysis depth. (D) Relative read frequency of protist ASVs having different habitats (EPI, EPI+MES, and MES) in each sample category (different depths and forms of 18S rRNA). ASVs with zero values in either measure were removed for all the plots.

The Sankey diagram shows an overall link between habitat depth and cell-lysis depth for protist ASVs (Fig. 6C). The majority (66.0%) of the ASVs significantly lysed in the epipelagic zone were originated from those ubiquitously distributed between the surface and deep (i.e., non-significant ASVs of the EEI measure), whereas 86.0% of the ASVs lysed in the mesopelagic zone were composed of epipelagic-specific ASVs.

The ASVs with a preference for epipelagic, mesopelagic, and both habitats comprised 67.9%, 1.4%, and 30.7% of the average relative contribution of cf-rRNA in the epipelagic zone, respectively (Fig. 6D). The average contribution of cf-rRNA in the mesopelagic zone was composed of ASVs in the epipelagic (7.8%), mesopelagic (33.6%), and both habitats (58.6 %).

## 4. Discussion

### 4.1. A quantitative assessment of intact and lysed eukaryotes

We established a simple method of extracting cell-free (pass through a 0.2 µm filter) RNA by directly capturing nucleic acids in a silica-membrane column. This method successfully recovered >80% of the rRNA dissolved in seawater over a wide concentration range, enabling quantitative and comprehensive analysis of dead and lysed microorganisms (Fig. 2A and 2B). This method demonstrated that over 90% of the cell-free 18S-rRNA was derived from protists (single-celled organisms such as phytoplankton and protozoa) rather than from multicellular organisms such as zooplankton and fishes. RNA is a highly labile molecule with a turnover time of a few hours to a few days due to its high bioavailability ^21^. Thus, cf-rRNA provides information on the taxonomy and degree of recent or ongoing cell lysis.

rRNA metabarcoding of the cellular fraction revealed that Prymnesiophyceae (Haptophyta) and Dinophyceae (Dinoflagellata) were the most active and ubiquitous phytoplankton lineages in the surface zone of the study sites (Fig. 3). However, Dinophyceae may include heterotrophic species, and their contributions may be overestimated in rRNA-based phytoplankton communities. Microscopic analysis further confirmed the dominance of haptophytes and dinoflagellates, with average contributions of 35.4% and 40.0%, respectively, to the total identifiable phytoplankton cell count (Fig. S3). Our results are consistent with previous observations showing that haptophytes are highly abundant (up to 1.01 × 10^4^ copies mL^−1^ of the 18S rRNA gene), contributing to 35–42% of the total chlorophyll *a* biomass in the Kuroshio ecosystem in autumn^34,35^.

### 4.2. Enhanced eukaryotic cell lysis in the mesopelagic zone

The concentration of cf-rRNA was much lower than that of ca-rRNA in the surface but comparably high in the mesopelagic zone, resulting in apparent increases in the cf-rRNA:ca-rRNA ratio in the mesopelagic zone (Fig. 2C). The low cf-rRNA:ca-rRNA ratio in the surface zone may be partly explained by the rapid consumption of dissolved ribosomes due to high bacterial abundance. However, a previous study demonstrated comparable rates of plankton-derived DOM consumption between surface and deep seawaters^36^. Thus, the release of cellular contents, including ribosomes, via eukaryotic cell lysis is likely enhanced in the mesopelagic layer. Increases in lytic activity and the subsequent remineralization of organic molecules in the mesopelagic zone have also been reported in previous studies in different environments ^14,37^.

This assertion is supported by the DA analysis results, which evaluated the degree of cell lysis for each protist ASV at each depth by comparing the rRNA composition between the living (cell-associated) and dissolved (cell-free) fractions. We detected a higher number of significantly lysed ASVs and a lower number of significantly intact (i.e., less susceptible to cell lysis) ASVs in the mesopelagic zone than in the surface ecosystems (Fig. 4). We also uncovered a distinct taxonomic signature in CLI measurements between epipelagic and mesopelagic zones, indicating depth-resolved differences in the cell-lysis dynamics of protists. Notably, diverse lineages represented not only by the protozoan Ciliophora but also by phytoplankton-including taxa Dinoflagellata and Haptophyta_X comprised significantly lysed ASVs in the mesopelagic layer (Fig. 4B). A comprehensive comparison revealed that both dominant and minor lineages, such as Choanoflagellata and Cryptophyta, were more susceptible to cell lysis in the mesopelagic layer than in the surface layer (Fig. 5A). These results consistently indicate that eukaryotic cell lysis was promoted below the sunlit epipelagic zone.

The lytic activity also differed within the mesopelagic zone, as indicated by higher cf-rRNA:ca-rRNA ratios in the upper (300 m) than in the middle mesopelagic layer (500 m; Fig. 2C). This might be attributed to the accumulation of sinking particles in the upper mesopelagic zone owing to the vertical density gradient of seawater. Approximately 1– 40% of photosynthetically fixed carbon is exported to the deep layer as cell aggregates or fecal pellets (biological carbon pump)^38^. However, a substantial fraction (ca. 30–90%) of the particle flux was shown to slow down or stop sinking in the upper mesopelagic zone, presumably because of the steep increase in seawater density^39^. The CTD profile showed steep density gradients of 100–300 m at the study sites (Fig. 2C). Therefore, organic particles exported from the surface layer might have been retained and intensively lysed in the upper mesopelagic zone.

### 4.3. Surface eukaryotes were lysed in the deep ocean

We showed that the mesopelagic zone (300–500 m) is a hotspot for viral lysis, and various lineages were actively lysed in this layer. These results prompt the question: What is the source of the protists intensively lysed in the deep sea? To address this, we investigated the relationship between preferred habitat depth and lysis measures for each ASV. Expectedly, heterotrophic lineages, such as Ciliophora, Radiolaria, and Choanoglagellata, were clustered into epipelagic-preferred and mesopelagic-preferred ASVs (Fig. 5C), indicating that these heterotrophic lineages are ubiquitous in the water column at the subdivision or class level but exhibit distinct habitat depths at the ASV level. Dinoflagellata and Cryptophyta also consist of surface- and deep-specific ASVs, likely reflecting their diverse trophic strategies (i.e., autotrophs, heterotrophs, and mixotrophs) within lineages^40^.

Notably, we found a significant positive correlation between cellular activity in the epipelagic and lytic mortality in mesopelagic zone across all microeukaryotes (Fig. 6A). Remarkably, most (86.0%) ASVs differentially (significantly) lysed in the mesopelagic phase were epipelagic-preferred species (Fig. 6B and 6C). Although they contributed 33.6% of the total cf-rRNA reads in the mesopelagic region, mesopelagic-preferred ASVs comprised only 1.8% of the differentially lysed ASVs (Fig. 6D). These results clearly indicate that a large majority of eukaryotic species lysed in the mesopelagic zone were not inhabiting it but were transported there from the upper surface layer via the biological carbon pump.

Although our study does not aim to provide conclusive evidence on the mechanisms underlying cell lysis, here we discuss two possible explanations for our results. First, virus-induced vertical export (“viral shuttle”) and subsequent cell lysis might explain cell sinking and destruction dynamics. Previous *in situ* observations showed that viral infection in the surface layer enhanced the sinking of infected cells of the haptophyte *Gephyrocapsa huxleyi*, which were subsequently lysed in the mesopelagic zone^14^. Consistently, plankton-infecting viruses in the mesopelagic zone were reported to be mainly transported from the surface zone with the sinking of infected cells^41^. A laboratory experiment also demonstrated that viral infection facilitated the aggregate formation of infected diatom cells, likely by producing proteinaceous extracellular polymers^42^. Considering the remarkable diversity and heterogeneity of eukaryotic viruses in marine environments^41,43,44^, “viral shuttle” and subsequent lysis at depth may be prevalent dynamics for various protist species, including heterotrophs.

The second possibility is that sloppy feeders such as copepods actively transformed the sinking particles to DOM. According to field observations, calanoid copepods consumed 84% of sinking organic carbon in the upper mesopelagic zone^45^. They would also have a role in mechanically degrading fast-sinking particles into slow-sinking particles^46^ that are subsequently lysed or demineralized in the mesopelagic zone. In the natural environment, viruses and sloppy feeders may contribute to cell lysis simultaneously and synergistically. Additional information on other biological factors, such as sloppy feeding, is required to gain mechanistic insights into the eukaryotic cell lysis.

The study area is influenced by the subtropical Kuroshio Current, characterized by low nutrient availability in the surface layer. Although oligotrophic regions typically have lower export fluxes than productive eutrophic regions, the transfer efficiency of biological carbon pumps is generally high in these regions, possibly due to low zooplankton degradation and high viral activity^38,47^. A recent observation in the same area reported that small phytoplankton and fecal pellets contributed to efficient carbon export^48^. Our results and previous findings underscore the importance of surface microeukaryotic communities in the biogeochemical cycles of oligotrophic oceans.

We revealed the importance of the mesopelagic zone in eukaryotic cell lysis and the epipelagic zone as the source of these cells using a newly established cell-free RNA analysis method. Dissolved RNA is a component of labile DOM that accounts for < 0.1% of the ocean DOC inventory yet 84% of the carbon turnover (flux of 15–25 Pg C yr^−1^)^49^. Viral lysis and sloppy feeding, which can be detected using our method, are estimated to contribute to approximately 45% of the labile DOC supply^49^, highlighting the need to unravel the source organisms and the underlying mortality mechanisms. Most labile DOC produced in the surface mixed layer is oxidized to CO_2_ via respiration and released into the atmosphere. In contrast, labile DOC released in the deep layer can be sequestered over a decadal timescale as dissolved inorganic carbon or refractory DOC because it is less susceptible to vertical mixing and photodegradation^49,50^. Thus, the export of plankton proliferating in the surface layer to the mesopelagic zone and their subsequent degradation may significantly impact the ocean carbon inventory on geological timescales. Ongoing ocean warming may modify the fate of fixed carbon in ocean biogeochemical cycles via shoaling the mixed layer depth, changes in the phytoplankton community, or decreases in viral infectivity^51,52^. Our findings provide a new perspective for evaluating the interplay between marine microorganisms and biogeochemical cycles in the ocean.

## Acknowledgement

We gratefully acknowledge the captain, officers, and crew of the R/V *Shinsei-Maru* for their support during the cruise. We also thank Hideki Fukuda for the arrangement of the cruise. This work was supported by JSPS KAKENHI (19H05667, 21H05057, 22H00384, 22H00385, 22H03716, 22H02420, and 23K23685), JST PRESTO (JPMJPR23G3), and JST CREST (JPMJCR23J4). Computational work was performed at the SuperComputer System, Institute for Chemical Research, Kyoto University.

## Data Accessibility and Benefit-Sharing

Raw sequencing reads are deposited in DDBJ Sequence Read Archive (DRA) under accession number DRA018638 (BioSample accessions SAMD00780584– SAMD00780643). Metadata is also stored in the DRA (PRJDB18111). Custom scripts and associated data files used in this study are available on GitHub (https://github.com/HisashiENDO/Kuroshio_cf-rRNA). Benefits Generated: Benefits from this research accrue from the sharing of our data and results on public databases as described above.

## Author contributions

H.E. designed this research. H.E. conducted the fieldwork and sampling. H.E., Y.Y., N.T. performed the laboratory experiments. H.E. performed the bioinformatics analysis. All the authors contributed to interpreting the data and writing the manuscript.

