## Supplementary file for "Extracellular rRNA profiling reveals the sinking and cell lysis dynamics of marine microeukaryotes"

**Table S1** Geographic locations of sampling sites.

| Station | Latitude (°N) | Longitude (°E) | Dampling date | Dampling time (JST) | Depth category | Depth (m) | Zone |
|---|---|---|---|---|---|---|---|
| 3 | 28.00 | 128.50 | 22-October 2022 | 0:38 | 10 m | 10 | Epipelagic |
| | | | | | SCM | 93 | Epipelagic |
| | | | | | MES | 300 | Mesopelagic |
| E1 | 28.50 | 128.39 | 23-October 2022 | 21:40 | 10 m | 10 | Epipelagic |
| | | | | | SCM | 54 | Epipelagic |
| | | | | | MES | 500 | Mesopelagic |
| 5 | 30.50 | 128.00 | 20-October 2022 | 23:45 | 10 m | 10 | Epipelagic |
| | | | | | SCM | 53 | Epipelagic |
| | | | | | MES | 300 | Mesopelagic |
| 6 | 30.63 | 128.55 | 21-October 2022 | 23:43 | 10 m | 10 | Epipelagic |
| | | | | | SCM | 35 | Epipelagic |
| | | | | | MES | 500 | Mesopelagic |
| 8 | 30.90 | 129.67 | 25-October 2022 | 9:22 | 10 m | 10 | Epipelagic |
| | | | | | SCM | 43 | Epipelagic |
| | | | | | MES | 300 | Mesopelagic |

**Table S2** Primer pairs used in this study.

| Experiments/Primer name | Sequence (5' to 3') | References |
|---|---|---|
| ***E. coli* strain B-specific 16S V4 primer** | | |
| 16S_EC442F | GCGGGGATGAAGGGAGTAAA | This study |
| 16S_EC636R | ATGCAGTTCCCAGGTTGAGC | This study |
| **Eukaryote-specific 18S V4 primer** | | |
| E572F | CYGCGGTAATTCCAGCTC | Comeau et al. (2011) |
| E1009R | AYGGTATCTRATCRTCTTYG | Comeau et al. (2011) |

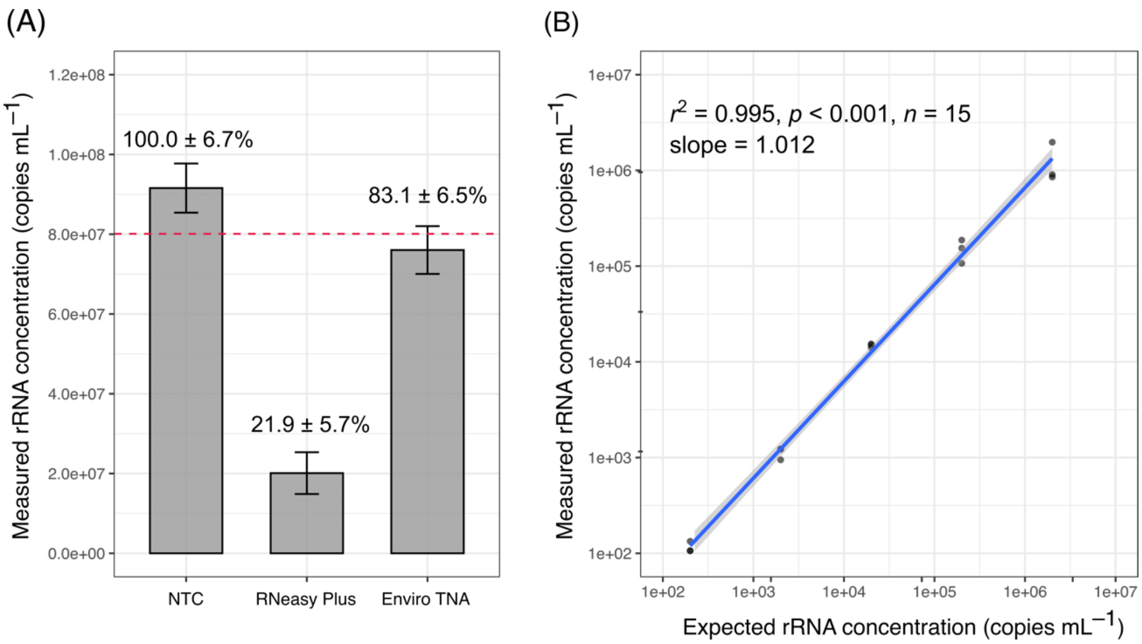

**Figure S1**. Performance of RNA extraction methods used. (A) Comparison of spiked-in rRNA concentrations (mean and SD, n = 3) recovered by two extraction kits and without extraction (i.e., non-treated control, NTC). The red dashed line denotes the expected values in the product data sheet. The value on top of each bar indicates the extraction efficiency calculated by comparing it with the NTC (mean and SD, n = 3). (B) Effect of spiked-in rRNA concentration on the extraction performance of the cell-free RNA extraction kit (Wizard Enviro TNA Kit). The linearity was evaluated in a 10-fold dilution series ranging from $2.0\times10^2$ to $2.0\times10^6$ copy $mL^{-1}$ (log-log linear regression: $r^2$ = 0.995, $p < 0.001$, $n$ = 15, the slope of 1.012 between the measured and expected values).

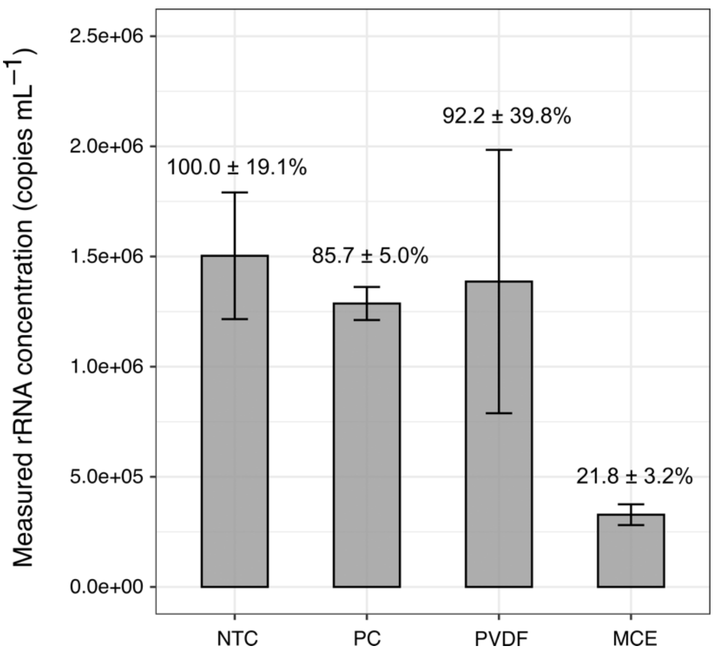

**Figure S2**. Performance of membrane filters used. Comparison of adsorption properties for ribosome (mean and SD, n = 3) with three filters made of polycarbonate (PC), polyvinylidene difluoride (PVDF), and mixed cellulose esters (MCE). The value on top of each bar indicates the recovery rate (mean and SD, n = 3) calculated by comparing it with the non-treated control (NTC).

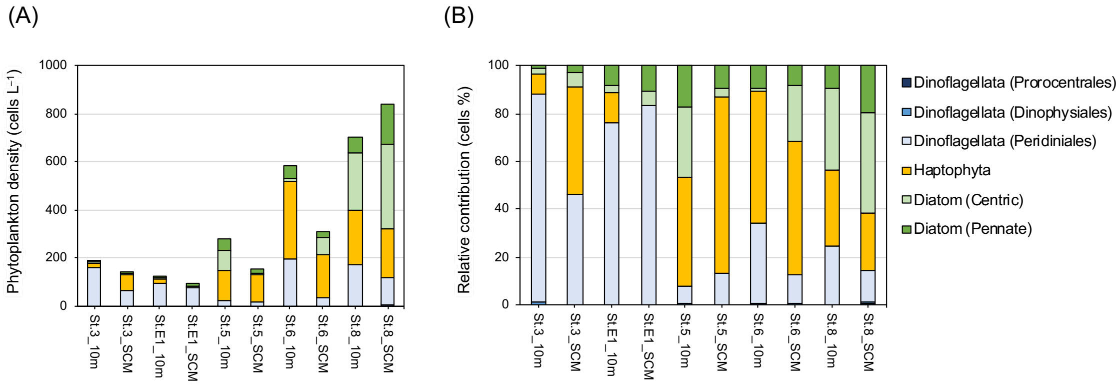

**Figure S3**. Phytoplankton cell counts were determined using light microscopy. (A) Abundance and (B) relative contributions of major phytoplankton in the epipelagic layer.
